# Dog brain atlas generated via spatially constrained spectral clustering

**DOI:** 10.1101/2025.11.25.690406

**Authors:** Raúl Hernández-Pérez, Laura V. Cuaya, Magdalena Boch, Fernando A. Barrios, Sarael Alcauter, Ludwig Huber, Claus Lamm

**Affiliations:** Social, Cognitive and Affective Neuroscience Unit, Department of Cognition, Emotion, and Methods in Psychology, Faculty of Psychology, University of Vienna, Vienna, Austria; Instituto de Neurobiología, Universidad Nacional Autónoma de México, Juriquilla, Querétaro, México; Messerli Research Institute, Department of Interdisciplinary Life Sciences, University of Veterinary Medicine Vienna, Veterinärplatz 1, 1210, Vienna, Austria

**Keywords:** dog brain atlas, spectral clustering, brain parcellation, rs-fMRI, functional connectivity, comparative neuroscience

## Abstract

Functional parcellations enable reproducible analyses of brain organization, yet the dog fMRI field still lacks a validated, multi-scale functional atlas. We applied spatially constrained spectral clustering to a resting state fMRI dataset (n = 27 dogs) to generate whole-brain parcellations spanning 20-300 parcels. Replicability was evaluated against an independent dataset (n = 20 dogs) using Dice overlap and Adjusted Rand Index (ARI) as measures of spatial overlap and similarity; within-parcel functional coherence was assessed via homogeneity. Functional parcellations were anatomically coherent across scales, with global Dice peaking at 0.625 at N = 140 and 0.623 at N = 100, while decreasing to 0.526 at N = 300. ARI peaked at 0.48 for N = 60 and remained ≥ 0.46 through N = 180. Mean within-parcel homogeneity increased monotonically from 0.08 (N = 20) to 0.19 (N = 300). Concordance with an anatomical atlas (Johnson et al., 2020) increased at the regional level (plateau ∼0.16 for N ≥ 140) while diminishing at gyral and lobar levels as resolution increased, consistent with functionally driven sub-regional differentiation. Together, these results indicate functional segmentations that are replicable across datasets and internally coherent across scales, with intermediate resolutions (100-140 parcels) balancing specificity and reproducibility for common analyses. We introduce a comprehensive, multi-scale functional dog brain atlas derived from data-driven clustering, providing an open resource for comparative studies of brain evolution and canine cognition.

## 1. Introduction

Mapping the brain into functionally coherent regions is essential for reproducible analyses of organization and functional connectivity (Craddock et al., 2012; Eickhoff et al., 2018) as well as for enabling comparisons across studies and species (Passingham et al., 2002). While standardized resources are widely available for human neuroimaging and anatomical atlases have been developed to support canine neuroimaging research (Czeibert et al., 2019; Datta et al., 2012; Johnson et al., 2020; Nitzsche et al., 2019), no existing resource yet provides functionally homogeneous parcels across multiple resolutions, leaving connectivity-driven studies of dog brain anatomy and function without a fit-for-purpose parcellation (Craddock et al., 2012; Eickhoff et al., 2018; Liu et al., 2018). In this study, we focused on five key properties that would make a functional dog atlas useful while at the same time fulfilling basic anatomical criteria: (i) spatial contiguity, i.e., parcels should be spatially connected to preserve anatomical interpretability (Thirion et al., 2006); (ii) multi-scale coverage, i.e., the atlas should provide parcellations at multiple levels of granularity, allowing users to match the scale to their specific research questions (Eickhoff et al., 2018; Parisot et al., 2016); (iii) reproducibility, i.e., parcellations must be robust and replicable across independent datasets and different acquisition protocols to ensure the reliability of the atlas (Rezende et al., 2019); (iv) internal coherence: each parcel should consist of functionally similar voxels, ensuring that the regions are functionally homogeneous (Craddock et al., 2012); (v) anatomical concordance, i.e., the functional parcels should align with established anatomical structures, which is essential for grounding functional findings within a known structural framework and relating function to the underlying anatomy (Eickhoff et al., 2018; Passingham et al., 2002).

Our aim is to provide a multi-scale functional atlas of the dog brain. To do so, we generated a functional dog atlas by applying a spatially constrained spectral clustering algorithm (Craddock et al., 2012; Wang et al., 2014) over different levels of granularity. Prior studies have demonstrated the suitability of spectral clustering methods for functional brain parcellation across different species and imaging modalities (Liu et al., 2018; Parisot et al., 2016; Xia et al., 2019). This approach assesses the five key properties as follows: (i) to ensure spatially contiguous parcels, it integrates a contiguity constraint; (ii) to achieve multi-scale coverage, we applied the algorithm at several levels of granularity (N = 20 to N = 300); (iii) to confirm reproducibility, we assessed both between-dataset agreement with an external dataset (Beckmann et al., 2021b) and within-participant robustness across different acquisition hardware (Guran et al., 2023) using the Dice coefficient and Adjusted Rand Index (ARI); (iv) we evaluated internal coherence using parcel homogeneity; and finally, (v) we examined anatomical concordance by comparing our functional segmentations against an established dog anatomical atlas (Johnson et al., 2020) at lobar, gyral, and regional levels.

We make this multi-resolution atlas publicly available to serve as a versatile tool for connectivity and task-based studies, facilitating comparative studies in neuroscience and contributing to a growing need for essential neuroimaging resources in canine as well as comparative canine-human neuroscience research (Boch et al., 2024; Bunford et al., 2017).

## 2. Methods

### 2.1 Participants

The study utilized two inhouse datasets, hereafter referred to as Inhouse K9 Dataset and the Inhouse Knee dataset.

The Inhouse K9 Dataset involved 27 pet dogs (12 females) of various breeds (ten Border Collies, five Australian Shepherds, three Labrador Retrievers, one Small Münsterländer, one Nova Scotia Duck Tolling Retriever, one Goldendoodle, and six mixed breeds) aged 2 to 12 years (mean = 5.96 years). Dogs were recruited from local pet owner communities and had been extensively trained to remain awake and motionless inside the MRI scanner following established training protocols (Karl et al., 2020).

The Inhouse Knee Dataset was acquired as part of the coil comparison study described in Guran et al. (2023). This dataset included 10 dogs (one Labrador Retriever, seven Border Collies, one Australian Shepherd, and one mixed breed) with an average age of 8.6 years. Every dog in this dataset is included in the K9 Dataset (Guran et al., 2023).

Caregivers of the participating dogs provided informed consent. All procedures were approved by the Ethics and Animal Welfare Commission of the University of Veterinary Medicine Vienna (No. ETK-06/06/2017) and conducted in accordance with national legislation, Good Scientific Practice, and the ARRIVE guidelines (Kilkenny et al., 2010). To assess the replicability and robustness of our findings, we utilized a publicly available external dataset “OpenNeuro Dataset ds003830” (Resting state fMRI of 17 idiopathic epileptic dogs and 20 healthy control dogs) by (Beckmann et al., 2021b) previously reported (Beckmann et al., 2020, 2021a). For our comparative analysis, we included only the healthy control group from the external dataset, which was comprised by 20 pet dogs

(ten female) of various breeds (ten Beagles, six Greater Swiss Mountain Dogs, and four Border Collies), aged 1 – 9 years (mean = 5.25 years). Inclusion in the control group required the dogs to be free of any history of seizures and to have passed normal clinical and neurological examinations, as well as a normal structural MRI of the brain. The dataset had been collected under a standardized sevoflurane anesthesia protocol with authorization from the Cantonal Veterinary Office of Zurich (approval numbers ZH272/16 and ZH046/20) and in compliance with the relevant guidelines and regulations.

### 2.2 Study design and animal welfare

The study followed the ARRIVE guidelines. Dogs were recruited consecutively based on owner availability and compliance with training protocols. No randomization was required because all individuals underwent identical imaging procedures; likewise, blinding was not feasible due to the nature of the image-acquisition workflow, though subsequent atlas construction and statistical analyses were performed using automated pipelines.

Inclusion criteria comprised the ability to complete awake MRI training, absence of neurological disease, and acceptable motion levels during scanning. Exclusion criteria were the presence of neurological abnormalities or excessive motion artifacts; no dogs were excluded.

### 2.3 Imaging Procedures

All imaging data for the inhouse K9 Dataset were collected using a 3T Siemens Skyra MR-system with an inhouse 16-channel receive coil (K9 coil; distributed by ALSIX GmbH, Austria) (Guran et al., 2023). Dogs wore earplugs secured with bandages to mitigate scanner noise and accessed the scanner via a custom-made ramp. Functional whole-brain images were acquired using a two-fold multiband-accelerated echo planar imaging (EPI) sequence. Data were obtained from 24 axial slices with interleaved acquisition in descending order and a 20% slice gap; repetition time (TR) = 1000 ms; echo time (TE) = 38 ms; flip angle = 61°; field of view (FOV) = 144×144×58 mm^3^; voxel size = 1.5×1.5×2 mm^3^). The average total acquisition time for the resting state scan was 7.6 minutes (range 6 – 8 minutes). Data for the Inhouse Knee Dataset were collected on the same 3T Siemens Skyra MR-system and with identical scan parameters; however, a standard human knee coil was used as the receive coil instead of the K9 coil (Guran et al., 2023). The total acquisition time for each scan was determined by the individual dog’s ability to remain motionless inside the scanner for a prolonged period. Shorter durations were utilized when a participant began to show signs of restlessness, ensuring that all collected data was of high quality and minimally affected by motion artifacts (Guran et al., 2023). The external dataset (Beckmann et al., 2020, 2021b, 2021a) had been acquired on a 3 Tesla Philips Ingenia scanner using a 16-channel dStream HeadSpine coil. In this dataset, dogs were anesthetized. Anesthesia was induced with propofol and maintained with sevoflurane, while key physiological parameters such as heart rate, blood pressure, and end-tidal CO₂ were continuously monitored and maintained within normal physiological ranges. Functional whole-brain images were acquired using an EPI sequence with the following parameters: TR = 2000 ms; TE = 30 ms; flip angle = 90°; FOV = 236 mm; slice thickness = 3 mm; voxel size = 1.5×1.5×3 mm^3^. The total acquisition time for the resting-state scan was approximately 12 minutes.

### 2.4 Data Analysis

The preprocessing of raw functional images of all three datasets was performed using FSL 6.0.5 (Jenkinson et al., 2012). We first applied motion correction and slice timing correction, then reoriented all the functional images to match the dog template (Nitzsche et al., 2019). We aligned all volumes and of the run and then calculated a mean functional image by averaging all volumes. We aligned the run with their corresponding mean functional image using FLIRT (Jenkinson et al., 2002). We then skull-stripped all images using a binary mask semi automatically drawn over the mean functional image. The mean functional image was transformed to the dog template using FLIRT, we used the resulting transformation matrix and applied it to each previously aligned run.

#### 2.4.1 Spatially constrained spectral clustering

We applied the method developed by (Craddock et al., 2012) called spatially constrained spectral clustering algorithm to all three datasets. This unsupervised machine learning method segments the brain by grouping voxels with similar functional MRI time-series, with the constraint that only spatially adjacent voxels can be clustered together. This ensures the resulting parcels are both functionally homogeneous and anatomically contiguous (Arslan et al., 2018). The clustering process was conducted iteratively, beginning with 20 parcels and increasing in steps of 20 up to 300 parcels. The use of the spatially constrained spectral clustering algorithm allowed us to generate segmentations ranging from coarse, large-scale networks to highly detailed functional sub-regions.

#### 2.4.2 Evaluation metrics

To evaluate the reliability of the generated dog brain segmentations using all three datasets and the anatomical atlas, we employed a series of metrics to quantify their spatial agreement and internal consistency. We used the Dice Coefficient to measure the spatial overlap, this metric measures the spatial overlap between segmentations. High Dice scores indicate strong alignment between segmentations of different datasets (Arslan et al., 2018). To assess the similarity between segmentations, we used the Adjusted Rand Index (ARI), this metric assesses the similarity between two segmentations, correcting for a possible by-chance agreement. An ARI score of 1 indicates perfect agreement between the segmentations, while a score of 0 indicates agreement equivalent to random chance. It quantifies the extent to which pairs of voxels are assigned to the same or different parcels in two segmentations, relative to what would be expected by random assignment (Arslan et al., 2018). Finally, the homogeneity metric was used to evaluate the internal consistency of each parcel (Craddock et al., 2012), evaluating the extent to which each parcel contains only voxels that belong to a single, coherent group (e.g., voxels with highly similar functional time-courses). A homogeneity score of 1 indicates that all parcels contain only data points which are members of a single class. It is a measure of the internal consistency of the parcels.

## 3. Results

The spectral clustering algorithm produced anatomically coherent parcellations across the full range of requested resolutions (N = 20-300; Figure 1A). At the coarsest level (20 parcels) the solution separated broad cortical and subcortical regions, while incrementally increasing N revealed progressively finer subdivisions that respected the gross boundaries of the coarse solution: large-scale borders are largely preserved as new parcels split from parent regions, indicating the expected hierarchical, nested organization resulting from the algorithm. This hierarchical property provides a multi-scale framework for analyzing brain organization from a global to a local level, capturing the spatial heterogeneity of brain architecture (Eickhoff et al., 2018; Gors et al., 2017).

**Figure 1.**
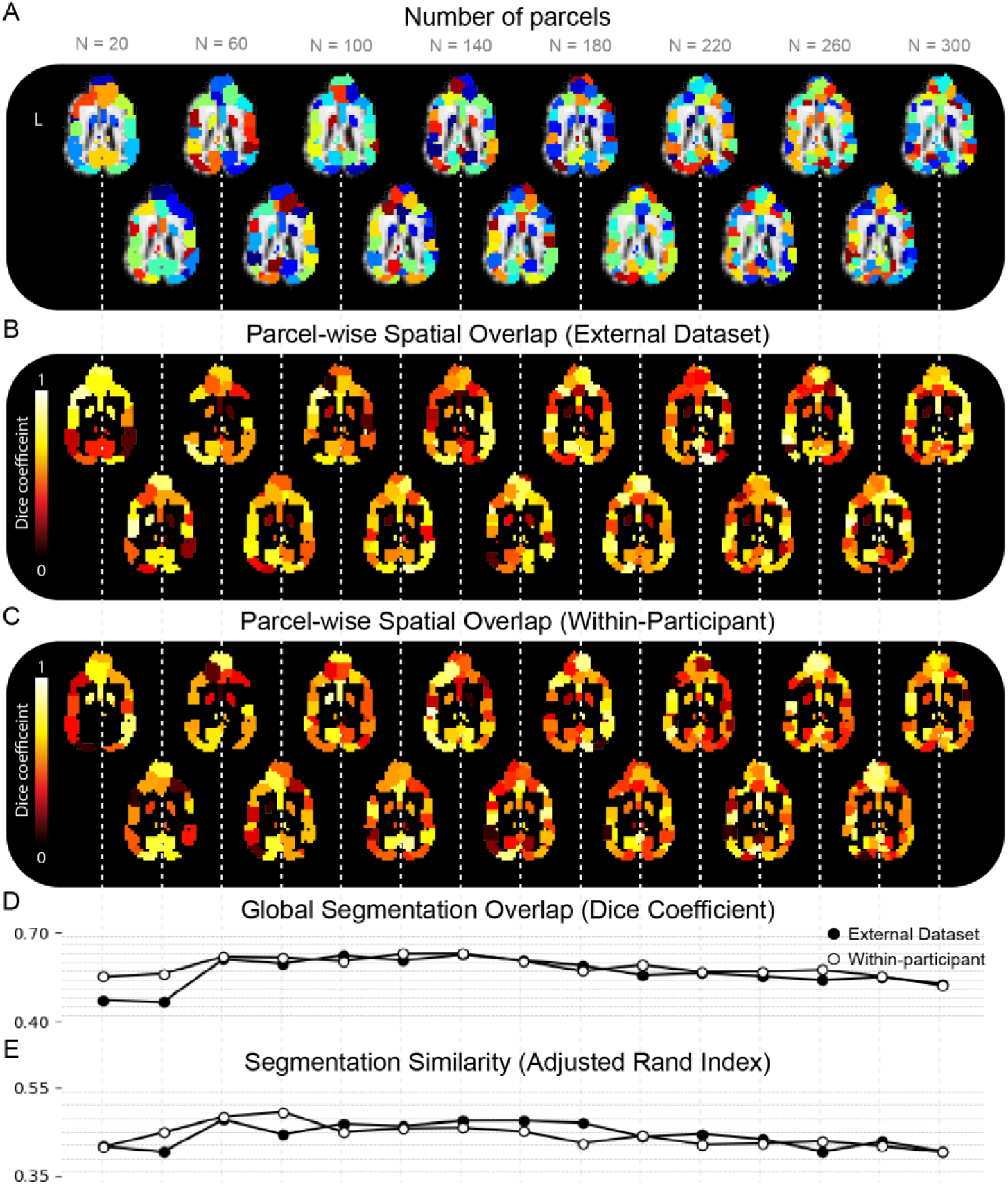
Brain segmentation using a spatially constrained spectral clustering algorithm and its cross-dataset replicability. **A.** Representative axial slice showing the dog brain at different levels of parcellation using a spatially constrained spectral clustering algorithm. The clustering algorithm was applied in steps of 20, from N=20 to 300 parcels, each column through the figure depicts the same slice at increasing levels of parcellation (see x-axis labels). Colors represent distinct parcels. **B.** Parcel-wise spatial overlap, the color indicates the Dice coefficient for individual parcels when compared to an external dataset (Beckmann et al., 2021b). **C.** Parcel-wise Overlap (Within-Participant): Parcel-wise spatial overlap (Dice coefficient) assessing within-participant robustness by comparing the Inhouse K9 Dataset (K9 coil) to the Inhouse Knee Dataset (human knee coil) (Guran et al., 2023). **D.** Global Overlap (Dice Coefficient): Global segmentation overlap, measured by the overall Dice coefficient for the entire parcellation at each resolution level, showing comparisons for both the external dataset and the within-participant dataset. **E.** Global Similarity (Adjusted Rand Index): Segmentation similarity, measured by the Adjusted Rand Index at each resolution level, showing comparisons for both the external dataset and the within-participant dataset.

### 3.1 Replicability against external dataset

To assess the consistency of individual parcels between the parcellations derived from our inhouse dataset and those from the independent external dataset (Beckmann et al., 2021b), we examined the distribution of Dice coefficients for all matched segment pairs at each level of parcellation (Fig. 1B). The results showed that despite the variability in global overlap, the median Dice coefficient for individual parcels remained high across all resolutions. At the 140-parcels parcellation, which demonstrated the highest global overlap, the median Dice coefficient for individual parcels was 0.58. Similarly high median values were observed for the 100-parcels (0.57) and 200-parcels (0.55) segmentations. This indicates a strong consistency at the level of individual parcels, suggesting that the spatially constrained spectral clustering algorithm is robust in identifying corresponding neuroanatomical units across the in-house dataset and the independent dataset (Beckmann et al., 2020, 2021b, 2021a).

Global overlap between the parcellations derived from our dataset and those generated with the external dataset was quantified with the Dice coefficient (Figure 1C). The results indicate an optimal range for the number of parcels. As shown in Figure 1C, segmentation accuracy was initially low for coarser parcellations. The coefficient increased significantly with the number of parcels, reaching a peak value of 0.625 at 140 parcels. A similarly high performance was observed for the 100-parcels segmentation, which showed a Dice coefficient of 0.623. Beyond this 100-140 parcels range, increasing the parcellation granularity resulted in a steady decline in segmentation accuracy, with the Dice coefficient decreasing to 0.526 for 300 parcels. This suggests that a segmentation of approximately 140 parcels provides the best compromise between spatial detail and inter-dataset reproducibility.

### 3.2 Within-Participant Replicability

To evaluate the replicability of the clustering solution we performed a within-participant comparison. This analysis determined the consistency of the clustering results for the same individuals across different imaging sessions and coil. The results indicate that replicability is optimized when using 140 clusters. At this level, the analysis yielded a maximum Dice coefficient of 0.629. The performance of the clustering solution showed a clear trend: the Dice coefficient generally improved as the number of clusters increased from 20 to 140. However, for solutions with more than 140 clusters, the Dice coefficient began to decrease, suggesting that a higher granularity did not lead to more stable results.

The Adjusted Rand Index (ARI) revealed a non-linear relationship between the number of parcels and the ARI score. The ARI was 0.42 at the coarsest segmentation level (N = 20) and reached its maximum value of 0.48 at N = 60 parcels. Following this peak, the ARI remained relatively stable and high for resolutions up to N = 180, with values consistently above 0.46 (e.g., 0.47 at N = 100, 0.47 at N = 140, 0.47 at N = 180). Beyond 180 parcels, the index showed a gradual decline, finishing at 0.41 for the finest-grained parcellation (N = 300). This pattern indicates substantial and robust agreement between the segmentations derived from the two independent datasets, particularly for parcel solutions between 60 and 180.

The ARI for the within-participant comparison showed a similar non-linear trend (Figure 1E, Within-participant). The ARI score was 0.42 at the 20-parcel level and reached its maximum value of 0.49 at N=80 parcels. Following this peak, the ARI score remained high (e.g., 0.483 at N=60, 0.457 at N=120, and 0.458 at N=140), before declining at finer scales after the 160-parcel level. This pattern confirms a high level of agreement and robustness in the parcellations, even when comparing data acquired with different receive coils (Guran et al., 2023).

The functional coherence within parcels was assessed by calculating the homogeneity of the generated brain parcels. The distribution of homogeneity values for individual segments is shown in Figure 2A: as the total number of parcels increases, the homogeneity of individual parcels also rises. This is an expected outcome, as smaller parcels are inherently more likely to group voxels with similar functional time-series, thus increasing their internal coherence (Arslan et al., 2018). The overall homogeneity of the parcels increased as the number of parcels grew (Figure 2B), while the mean homogeneity, averaged across all parcels, showed a steady increase from approximately 0.08 at a 20-parcel segmentation to 0.19 at a 300-parcel segmentation. This finding aligns with previous work demonstrating that a higher number of parcels in data-driven segmentations leads to increased functional homogeneity, reflecting a trade-off between the level of detail and broader anatomical interpretability (Craddock et al., 2012).

**Figure 2.**
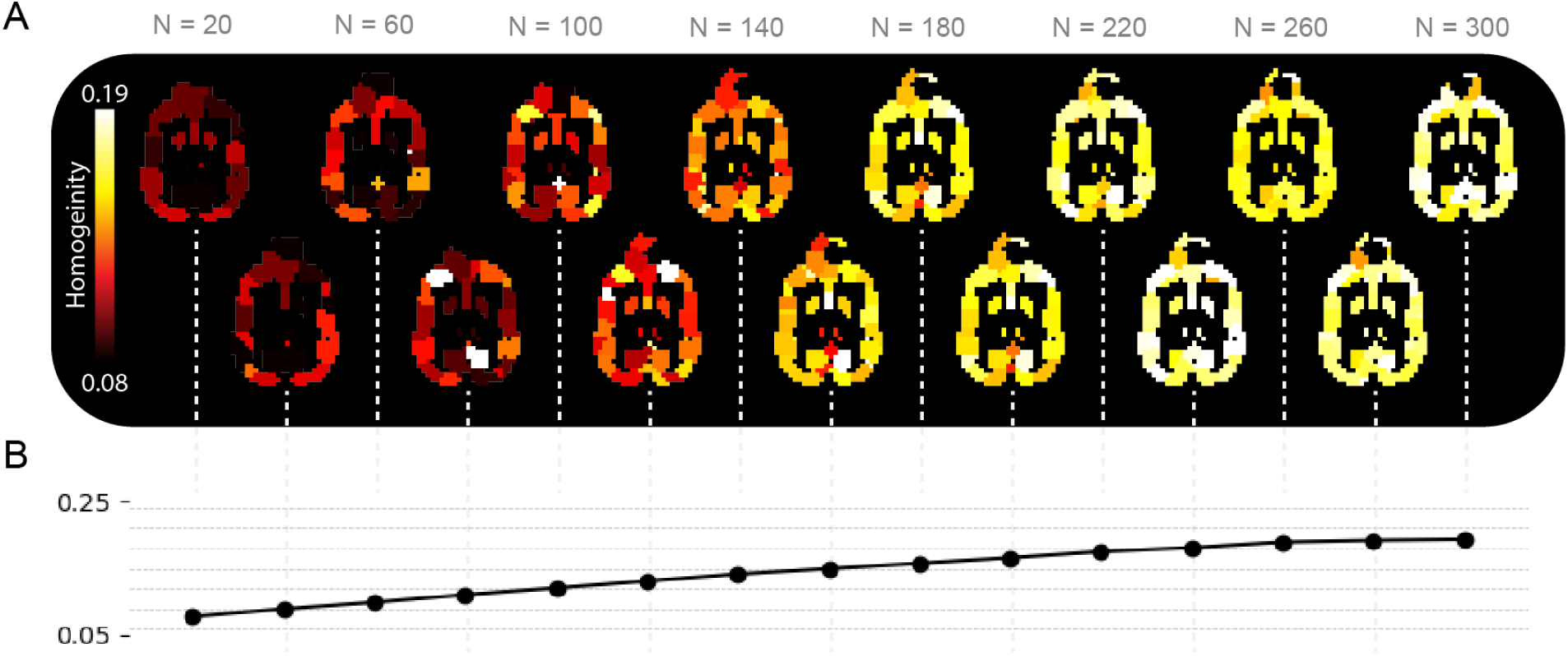
Within-cluster homogeneity across multi-scale brain parcellations. **A.** Homogeneity visualized on the same representative axial slice shown in Figure 1; For every parcellation level, lighter colors (yellow-white) denote higher within-cluster homogeneity whereas darker colors (deep red) indicate lower values (color bar, 0.08 - 0.19). **B.** Average homogeneity index obtained for each parcellation level produced by the spatially-constrained spectral clustering algorithm (N = 20 - 300). Higher values indicate that voxels grouped within the same cluster share more similar time-courses.

### 3.3 Concordance with the anatomical atlas

The similarity between the data-driven segmentations and the Johnson atlas (Johnson et al., 2020) varied systematically on both the number of clusters requested by the algorithm and the anatomical granularity of the atlas (Figure 3). Across the 20 - 300 parcel ranges the Dice coefficient displayed changes that closely corresponded to the atlas hierarchy (Fig. 3A). The dice coefficient decreased as the number of parcels increased at both the lobar and the gyral level. However, at the regional level, the dice coefficient demonstrated an inverse trend, increasing from approximately 0.02 at N = 20 to a plateau of roughly 0.16 for segmentations with 140 or more parcels. The segment-wise dice coefficients shown in Figure 3B-D confirm these results, showing intensifying regional overlap and diminishing gyral and lobar correspondence as the number of segments increases. This suggests that as the parcellation becomes finer, the algorithm-defined clusters show greater spatial correspondence with established anatomical regions.

**Figure 3.**
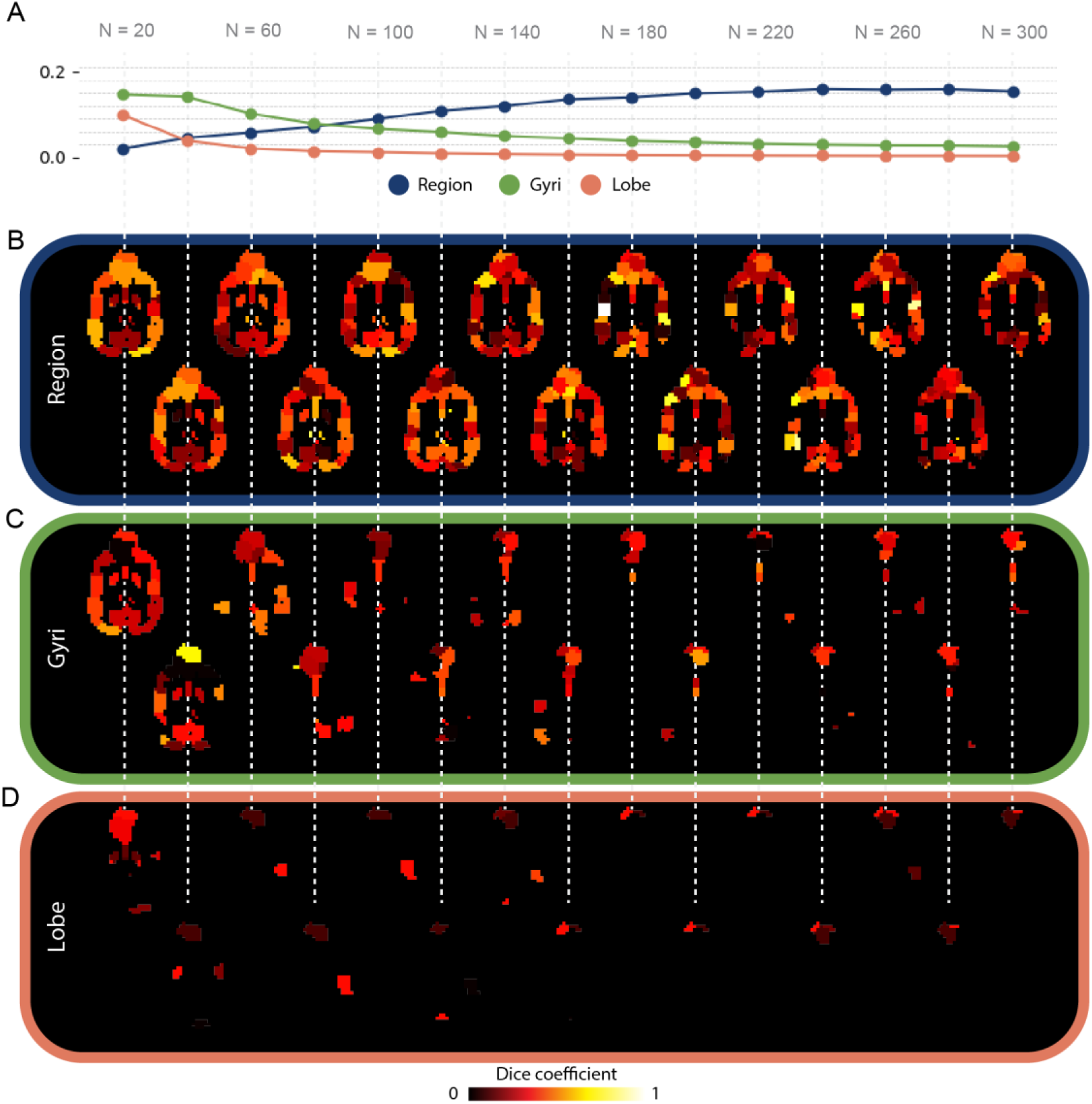
Concordance between the data-driven parcellations and the anatomical atlas (Johnson et al., 2020). **A.** Dice-coefficient for every parcellation level (N = 20 - 300 clusters; x-axis) for atlas regions (blue), gyri (green) and lobes (orange). B–D. Voxel-wise Dice maps for the same resolutions: region level (**B**), gyrus level (**C**) and lobe level (**D**). The color bar represents the Dice coefficient, with values ranging from 0 (no spatial overlap) to 1 (perfect spatial overlap). Region overlap increases as parcels become finer, whereas gyrus and lobe overlap decline.

## 4. Discussion

We generated a multi-scale functional dog brain atlas using spatially constrained spectral clustering on an awake resting state fMRI dataset of 27 dogs, producing parcellations from 20 to 300 clusters. The atlas shows robust replicability, both in a within-participant comparison across different acquisition hardware (Guran et al., 2023) and in a cross-dataset agreement test with an independent dataset (Beckmann et al., 2021b; N = 20). Global Dice coefficients, as a measure of parcel spatial congruency, peaked at 0.629 (N=140) for the within-participant test and 0.625 (N=140) for the external dataset, declining modestly at finer scales (0.526 at N = 300 for the external test). The Adjusted Rand Index (ARI), as a measure of parcel similarity, showed high stability, peaking at 0.49 (N=80) for the within-participant test and 0.48 (N=60) for the external dataset. ARI values remained high (≥0.46 for the external test and ≥0.45 for the within-participant test) across a broad range of intermediate resolutions. Within-parcel functional homogeneity increased regularly with granularity (0.08 at N = 20 to 0.19 at N = 300), as expected for a functional atlas. Together, these metrics indicate parcellations that are both replicable and internally coherent across scales.

Concordance with an anatomical atlas (Johnson et al., 2020) varied by hierarchy. As the number of parcellations increased, regional-level overlap increased (plateau ∼0.16 at N≥140), while gyral/lobar overlap decreased, consistent with functionally driven parcellations refining sub-regional distinctions as resolution increases. This divergence is expected, as anatomical homogeneity does not guarantee functional homogeneity (Craddock et al., 2012). The function of any cortical area is largely determined by its unique pattern of connections, or its connectional fingerprint (Passingham et al., 2002).

Consequently, a single anatomically defined gyrus might contain several functionally distinct modules. As our parcellation becomes more fine-grained, the algorithm identifies these smaller, more functionally coherent units, which often refine the underlying anatomy by subdividing or crossing traditional anatomical borders. This finding aligns with the principle that different parcellation strategies capture unique and valid aspects of brain organization; our data-driven approach reveals a map based on functional connectivity rather than macroscopic anatomical landmarks (Eickhoff et al., 2018).

Using an unsupervised, data-driven method avoids reliance on manual delineations (which are time-consuming and variable across individuals) or single-atlas priors (limited functional coherence) and instead derives parcels directly from intrinsic connectivity structure (Craddock et al., 2012; González-Villà et al., 2016). The observed cross-dataset reproducibility indicates that spectral clustering extracts stable functional divisions even when the data source changes. This is a critical advantage, as functional areas do not always align neatly with anatomical boundaries; therefore, a parcellation derived directly from functional data is more likely to represent functionally coherent units (Arslan et al., 2018; Eickhoff et al., 2018). Our results, which demonstrate increasing homogeneity with a higher number of parcels (Figure 2), are consistent with findings from human neuroimaging, which also show a trade-off between the interpretability of larger regions and the functional specificity of smaller, more numerous parcels (Craddock et al., 2012). The ability to generate consistent parcellations across independent datasets is a crucial step toward establishing a reliable and standardized functional atlas for the canine brain, addressing a significant gap in comparative neuroscience.

The pattern of increasing within-parcel homogeneity with granularity, alongside stable ARI over a broad most numbers of parcels, suggests that intermediate resolutions (100-140 parcels) strike the best balance between functional specificity and inter-dataset reproducibility for many analytic contexts. This finding aligns with previous human neuroimaging research, which established that while finer-grained parcellations increase functional homogeneity, the number of parcels is the most critical factor for creating brain atlases suited for functional connectivity analysis (Craddock et al., 2012). Finer parcellations (more clusters) naturally produce more homogeneous regions, but they can also capture noise specific to the dataset, potentially reducing generalizability. Our results reflect this trade-off: homogeneity steadily increases with the number of clusters, while the Dice coefficient and ARI plateau between approximately 100 and 180 parcels before showing a slight decline, indicating a point of diminishing returns where increased detail may not improve cross-dataset replicability. The choice of parcellation method and, crucially, the number of parcels, can significantly impact the results of network analyses (Arslan et al., 2018). Therefore, by providing a multi-resolution atlas, we offer a resource that allows researchers to select the level of granularity that best fits their specific research question, balancing the need for functional detail with robust, reproducible findings.

Our atlas provides a functional, multi-scale resource that fills an important gap in currently available tools in the dog neuroimaging field. While existing resources, such as anatomical templates, white-matter atlases and structural connectivity maps (Cordeau et al., 2025; Czeibert et al., 2019; Datta et al., 2012; Inglis et al., 2024; Johnson et al., 2020; Nitzsche et al., 2019), offer essential structural references, they do not inherently delineate the brain’s functional organization in all regions, which is known, subdivide, or cross these established anatomical borders. By applying spatially constrained spectral clustering, a method validated across human and comparative neuroimaging studies (Craddock et al., 2012; Parisot et al., 2016), we introduce a data-driven atlas that reflects intrinsic functional architecture while preserving anatomical contiguity. This makes our atlas particularly suitable for dog functional connectivity analyses.

An important finding of our study is the high degree of replicability demonstrated in two separate validation tests: the segmentations generated from our awake-dog dataset (using a 15-channel coil) showed high consistency both with (1) those from an external dataset acquired under sevoflurane anesthesia (Beckmann et al., 2021b), and (2) those generated from the same participants using different acquisition hardware (a human knee coil) (Guran et al., 2023). The consistency with the anesthetized dataset can be attributed to the fact that our segmentation approach, spatially constrained spectral clustering, operates on intrinsic functional connectivity patterns. The large-scale functional architecture defined by these patterns has been shown to remain largely preserved under light anesthesia, such as the one used in the reference dataset, with studies demonstrating that resting-state networks in dogs under sevoflurane are similar to those found in awake dogs (Beckmann et al., 2020). Furthermore, the high agreement found in the within-participant hardware comparison (Guran et al., 2023) demonstrates the robustness of the parcellations to variations in the acquisition setup. Spatially constrained spectral clustering, therefore, appears to successfully capture these stable, underlying network structures that persist despite differing states of consciousness and acquisition hardware, allowing it to generate robust and comparable parcellations across both datasets.

Despite our approach holds great promise and is based on robust and validated methodology, some limitations of this study should be acknowledged. First, the significant morphological diversity across dog breeds, including differences in skull types (Barton et al., 2024; Czeibert et al., 2020) and breed-related brain specializations (Hecht et al., 2019), presents a persistent challenge for creating a universally applicable brain atlas. While our study utilized a breed-averaged template (Nitzsche et al., 2019) to mitigate this issue, future research involving larger and more diverse dog breeds is necessary to refine these parcellations for specific cranial conformations. Second, our atlas is derived exclusively from resting-state fMRI data using a single parcellation algorithm. The integration of multi-modal data, such as structural MRI and diffusion tensor imaging, could yield a more comprehensive and biologically robust atlas, as different imaging markers capture unique aspects of brain organization (Eickhoff et al., 2018). A systematic comparison with alternative parcellation methods would also be beneficial for a complete assessment of the strengths and weaknesses of the spectral clustering approach for the dog brain (Arslan et al., 2018). Finally, our validation was based on computational metrics of replicability (against an external fMRI dataset) and anatomical concordance against the Johnson et al. (2020) atlas. While the Johnson et al. atlas is itself derived from histological maps, a direct, subject-specific histological validation of our functional parcels was not feasible, which remains a fundamental challenge in the field.

In conclusion, we introduce a reproducible, multi-resolution functional atlas of the dog brain derived from data-driven clustering, and we make it open access available on https://doi.org/10.5281/zenodo.17559991. We hope this resource will support standardized functional connectivity analyses, enable cross-study comparisons, and establishes a strong foundation for future multimodal and comparative atlas-building efforts.

## Data and Code Availability

The atlas is publicly available at https://doi.org/10.5281/zenodo.17559991. The code supporting this manuscript can be accessed at the following GitHub repository: https://github.com/rhernandez00/spatially_constrained_spectral_clustering.

## Author Contributions

R.H.P.: conceptualization, data curation, formal analysis, methodology, project administration, software, validation, visualization, writing – original draft, writing – review & editing. L.V.C.: conceptualization, writing – original draft, writing – review & editing. M.B.: data curation, investigation, writing – review & editing. F.A.B: methodology, writing – review & editing. S.A.: methodology, writing – review & editing. L.H.: funding acquisition, resources, supervision, writing – review & editing. C.L.: conceptualization, funding acquisition, resources, supervision, writing – review & editing.

## Acknowledgements

We are grateful to all the dogs and their caregivers for their participation in this project. We thank Laura Lausegger, Marion Umek, and Sabrina Karl for their dedication in training the dogs and assisting with data collection, together with Anna Thallinger, Katharina Fiala, Sara Binder, Olaf Borghi, and the many interns who contributed to data acquisition at the Comparative Canine Neuroimaging Unit. We also thank Erik Pasaye for his technical support. We acknowledge Katrin M. Beckmann (Neurology Department, Clinic of Small Animal Surgery, Vetsuisse Faculty Zurich) and her collaborators for making their dog rs-fMRI dataset publicly available. And finally, we thank R. Cameron Craddock and collaborators for developing and openly sharing the code for the spatially constrained spectral clustering algorithm that made this work possible.

## References

Arslan, S., Ktena, S. I., Makropoulos, A., Robinson, E. C., Rueckert, D., & Parisot, S. (2018). Human brain mapping: A systematic comparison of parcellation methods for the human cerebral cortex. NeuroImage, 170, 5–30. 10.1016/j.neuroimage.2017.04.014

Barton, S. A., Kent, M., & Hecht, E. E. (2024). Covariation of Skull and Brain Morphology in Domestic Dogs. Journal of Comparative Neurology, 532(9), e25668. 10.1002/cne.25668

Beckmann, K. M., Wang-Leandro, A., Dennler, M., Carrera, I., Richter, H., Bektas, R. N., Steiner, A., & Haller, S. (2020). Resting state networks of the canine brain under sevoflurane anaesthesia. PLOS ONE, 15(4), e0231955. 10.1371/journal.pone.0231955

Beckmann, K. M., Wang-Leandro, A., Richter, H., Bektas, R. N., Steffen, F., Dennler, M., Carrera, I., & Haller, S. (2021a). Increased resting state connectivity in the anterior default mode network of idiopathic epileptic dogs. Scientific Reports, 11(1), 23854. 10.1038/s41598-021-03349-x

Beckmann, K. M., Wang-Leandro, A., Richter, H., Bektas, R. N., Steffen, F., Dennler, M., Carrera, I., & Haller, S. (2021b). Resting state fMRI of 17 idiopathic epileptic dogs and 20 healthy control dogs (Version 1.0.0) [Dataset]. OpenNeuro. 10.18112/OPENNEURO.DS003830.V1.0.0

Boch, M., Huber, L., & Lamm, C. (2024). Domestic dogs as a comparative model for social neuroscience: Advances and challenges. Neuroscience & Biobehavioral Reviews, 162, 105700. 10.1016/j.neubiorev.2024.105700

Bunford, N., Andics, A., Kis, A., Miklósi, Á., & Gácsi, M. (2017). Canis familiaris As a Model for Non-Invasive Comparative Neuroscience. Trends in Neurosciences, 40(7), 438–452. 10.1016/j.tins.2017.05.003

Cordeau, M., Barton, S. A., & Hecht, E. E. (2025). Organization of distributed cortical connections underlying the processing of auditory information in dogs assessed by diffusion MRI. Imaging Neuroscience, 3, IMAG.a.16. 10.1162/IMAG.a.16

Craddock, R. C., James, G. A., Holtzheimer, P. E., Hu, X. P., & Mayberg, H. S. (2012). A whole brain fMRI atlas generated via spatially constrained spectral clustering. Human Brain Mapping, 33(8), 1914–1928. 10.1002/hbm.21333

Czeibert, K., Andics, A., Petneházy, Ö., & Kubinyi, E. (2019). A detailed canine brain label map for neuroimaging analysis. Biologia Futura, 70(2), 112–120. 10.1556/019.70.2019.14

Czeibert, K., Sommese, A., Petneházy, Ö., Csörgő, T., & Kubinyi, E. (2020). Digital Endocasting in Comparative Canine Brain Morphology. Frontiers in Veterinary Science, 7, 565315. 10.3389/fvets.2020.565315

Datta, R., Lee, J., Duda, J., Avants, B. B., Vite, C. H., Tseng, B., Gee, J. C., Aguirre, G. D., & Aguirre, G. K. (2012). A Digital Atlas of the Dog Brain. PLoS ONE, 7(12), e52140. 10.1371/journal.pone.0052140

Eickhoff, S. B., Yeo, B. T. T., & Genon, S. (2018). Imaging-based parcellations of the human brain. Nature Reviews Neuroscience, 19(11), 672–686. 10.1038/s41583-018-0071-7

González-Villà, S., Oliver, A., Valverde, S., Wang, L., Zwiggelaar, R., & Lladó, X. (2016). A review on brain structures segmentation in magnetic resonance imaging. Artificial Intelligence in Medicine, 73, 45–69. 10.1016/j.artmed.2016.09.001

Gors, D., Suetens, P., Vandenberghe, R., & Claes, P. (2017). Hierarchical spectral clustering of MRI for global-to-local shape analysis: Applied to brain variations in Alzheimer’s disease. 2017 IEEE 14th International Symposium on Biomedical Imaging (ISBI 2017), 787–791. 10.1109/ISBI.2017.7950636

Guran, C.-N. A., Sladky, R., Karl, S., Boch, M., Laistler, E., Windischberger, C., Huber, L., & Lamm, C. (2023). Validation of a New Coil Array Tailored for Dog Functional Magnetic Resonance Imaging Studies. Eneuro, 10(3), ENEURO.0083-22.2022. 10.1523/ENEURO.0083-22.2022

Hecht, E. E., Smaers, J. B., Dunn, W. D., Kent, M., Preuss, T. M., & Gutman, D. A. (2019). Significant Neuroanatomical Variation Among Domestic Dog Breeds. The Journal of Neuroscience, 39(39), 7748–7758. 10.1523/JNEUROSCI.0303-19.2019

Inglis, F. M., Taylor, P. A., Andrews, E. F., Pascalau, R., Voss, H. U., Glen, D. R., & Johnson, P. J. (2024). A diffusion tensor imaging white matter atlas of the domestic canine brain. Imaging Neuroscience, 2, imag–2–00276. 10.1162/imag_a_00276

Jenkinson, M., Bannister, P., Brady, M., & Smith, S. (2002). Improved Optimization for the Robust and Accurate Linear Registration and Motion Correction of Brain Images. NeuroImage, 17(2), 825–841. 10.1006/nimg.2002.1132

Jenkinson, M., Beckmann, C. F., Behrens, T. E. J., Woolrich, M. W., & Smith, S. M. (2012). FSL. NeuroImage, 62(2), 782–790. 10.1016/j.neuroimage.2011.09.015

Johnson, P. J., Luh, W.-M., Rivard, B. C., Graham, K. L., White, A., FitzMaurice, M., Loftus, J. P., & Barry, E. F. (2020). Stereotatic Cortical Atlas of the Domestic Canine Brain. Scientific Reports, 10(1), 4781. 10.1038/s41598-020-61665-0

Karl, S., Boch, M., Zamansky, A., Van Der Linden, D., Wagner, I. C., Völter, C. J., Lamm, C., & Huber, L. (2020). Exploring the dog–human relationship by combining fMRI, eye-tracking and behavioural measures. Scientific Reports, 10(1), 22273. 10.1038/s41598-020-79247-5

Kilkenny, C., Browne, W., Cuthill, I. C., Emerson, M., & Altman, D. G. (2010). Animal research: Reporting *in vivo* experiments: The ARRIVE guidelines. British Journal of Pharmacology, 160(7), 1577–1579. 10.1111/j.1476-5381.2010.00872.x

Liu, C., Ye, F. Q., Yen, C. C.-C., Newman, J. D., Glen, D., Leopold, D. A., & Silva, A. C. (2018). A digital 3D atlas of the marmoset brain based on multi-modal MRI. NeuroImage, 169, 106–116. 10.1016/j.neuroimage.2017.12.004

Nitzsche, B., Boltze, J., Ludewig, E., Flegel, T., Schmidt, M. J., Seeger, J., Barthel, H., Brooks, O. W., Gounis, M. J., Stoffel, M. H., & Schulze, S. (2019). A stereotaxic breed-averaged, symmetric T2w canine brain atlas including detailed morphological and volumetrical data sets. NeuroImage, 187, 93–103. 10.1016/j.neuroimage.2018.01.066

Parisot, S., Arslan, S., Passerat-Palmbach, J., Wells, W. M., & Rueckert, D. (2016). Group-wise parcellation of the cortex through multi-scale spectral clustering. NeuroImage, 136, 68–83. 10.1016/j.neuroimage.2016.05.035

Passingham, R. E., Stephan, K. E., & Kötter, R. (2002). The anatomical basis of functional localization in the cortex. Nature Reviews Neuroscience, 3(8), 606–616. 10.1038/nrn893

Rezende, T. J. R., Campos, B. M., Hsu, J., Li, Y., Ceritoglu, C., Kutten, K., França Junior, M. C., Mori, S., Miller, M. I., & Faria, A. V. (2019). Test–retest reproducibility of a multi-atlas automated segmentation tool on multimodality brain MRI. Brain and Behavior, 9(10), e01363. 10.1002/brb3.1363

Thirion, B., Flandin, G., Pinel, P., Roche, A., Ciuciu, P., & Poline, J. (2006). Dealing with the shortcomings of spatial normalization: Multi-subject parcellation of fMRI datasets. Human Brain Mapping, 27(8), 678–693. 10.1002/hbm.20210

Wang, X., Qian, B., & Davidson, I. (2014). On constrained spectral clustering and its applications. Data Mining and Knowledge Discovery, 28(1), 1–30. 10.1007/s10618-012-0291-9

Xia, X., Fan, L., Hou, B., Zhang, B., Zhang, D., Cheng, C., Deng, H., Dong, Y., Zhao, X., Li, H., & Jiang, T. (2019). Fine-Grained Parcellation of the Macaque Nucleus Accumbens by High-Resolution Diffusion Tensor Tractography. Frontiers in Neuroscience, 13, 709. 10.3389/fnins.2019.00709

